# DPGT: A spark based high-performance joint variant calling tool for large cohort sequencing

**DOI:** 10.64898/2026.03.02.709184

**Authors:** Chun Gong, Qi Yang, Ruiwen Wan, Shengkang Li, Yong Zhang, Yuxiang Li

**Affiliations:** BGI Research, Wuhan 430074, China; State Key Laboratory of Genome and Multi-omics Technologies, BGI Research, Shenzhen 518083, China

**Keywords:** SNP/INDEL, joint calling, computational performance optimization, parallel computing

## Abstract

**Background:** Joint variant calling is a crucial step in population-scale sequencing analysis. While population-scale sequencing is a powerful tool for genetic studies, achieving fast and accurate joint variant calling on large cohorts remains computationally challenging.

**Findings:** To meet this challenge, we developed Distributed Population Genetics Tool (DPGT), an efficient computing framework and a robust tool for joint variant calling on large cohorts based on Apache Spark. DPGT simplifies joint calling tasks for large cohorts with a single command on a local computer or a computing cluster, eliminating the need for users to create complex parallel workflows. We evaluated the performance of DPGT using 2,504 1000 Genomes Project (1KGP), 6 Genome in a Bottle (GIAB) and 9,158 internal whole genome sequencing (WGS) samples together with existing methods. As a result, DPGT produced results comparable in accuracy to existing methods, with less time and better scalability.

**Conclusions:** DPGT is a fast, scalable, and accurate tool for joint variant calling. The source code is available under a GPLv3 license at https://github.com/BGI-flexlab/DPGT, implemented in Java and C++.

## Introduction

With the development of sequencing technology, large cohort studies are increasingly available, such as the Genome Aggregation Database(gnomAD; 76,156 WGS samples)[1], Trans-Omics for Precision Medicine (TOPMed; ∼180,000 samples)[2], UK Biobank(UKBB; half a million UK participants)[3] ranging in sample size from tens of thousands to half a million. Large cohort sequencing deepens our understanding of how genetics influences disease[4,5], thereby facilitating advancements in precision medicine.

Joint calling is a method that combines individual variants into a population variant matrix, which improves variant discovery sensitivity and is an essential step for cohort sequencing analysis. Assembly-based variant callers such as Platypus and Genome Analysis Toolkit(GATK) HaplotypeCaller can perform joint calling with high INDEL accuracy by processing alignment files(BAMs) from multiple samples. However, their scalability is limited, as graph complexity and memory usage increase exponentially with sample size.[6]. To address this, the GATK team developed Reference Confidence Model (RCM) and genomic variant call format(gVCF) for joint calling on large scale samples[6]. The RCM estimates the likelihoods over the set of unobserved, non-reference alleles and genotypes, facilitating joint genotyping. Many widely used variant callers, including GATK HaplotypeCaller[6], DeepVariant[7], Sentieon HaplotypeCaller[8], Illumina DRAGEN[9], Streka2[10] and freebayes[11], can output variants in gVCF format. Both GATK (GenomicsDBImport and GenotypeGVCFs) and GLnexus[12] utilize gVCF files for joint calling, achieving improved scalability.

However, joint calling is compute-intensive and time consuming for large cohorts[3]. Using GATK for joint calling of a large number of samples faces issues with I/O, memory and scalability. For example, joint calling on 150K UKB WGS samples using GATK took 9.6 million CPU hours, with each CPU reserved 16.6GB of RAM. Under the condition of providing 1,458GB of memory, there were still 320 tasks failed due to insufficient memory or program errors. Moreover, we observed that the time for GATK to calculate the maximum likelihood expectation (MLE) of the allele counts (MLEAC) and the allele frequency (MLEAF) increases significantly with an increase in the number of alleles (AC), which limits the ability to scale effectively to larger sample size. The publicly available open-source GLnexus is an efficient joint caller, however, it lacks features for production use[12]. It does not support computing cluster and does not support randomly fetching variants from a target region from variant call format (VCF) files with the vcf index file provided. Running GLnexus in parallel on a computing cluster requires users to pre-extract variants from target regions, which will increase I/O and runtime. Here we present DPGT, a distributed joint calling tool for large cohorts based on Apache Spark. We benchmarked our tool against open-source tools GATK[6,13] and GLnexus[12,14] using 2,510 whole genome sequencing(WGS) samples(including 2,504 samples from 1000 Genomes Project(1KGP)[15] and 6 samples from Genome in a Bottle(GIAB)), 9,158 internal WGS samples and 100,000 simulated samples. DPGT produced results comparable in accuracy to existing methods, with less runtime and better scalability. Furthermore, DPGT has several useful features for production: it supports computing clusters and can resume tasks after interruption (Supplementary Table S1).

### Findings

#### Joint calling workflow

The workflow of DPGT is presented in Figure 1. Briefly, DPGT parallelizes the joint calling process by partitioning on two dimensions: the sample dimension and the genome position dimension. The input of DPGT is a list of gVCF files, which can be generated by GATK[6,13], Sentieon[8] or Illumina DRAGEN[9], and a reference fasta file. First, DPGT combines gVCF headers of all samples. Next, the whole genome region is divided into many non-overlapping windows. In each window, DPGT performs variant sites finding, variants combining and genotyping. Finally, DPGT concatenates genotyped VCF files into one result VCF file.

**Figure 1.**
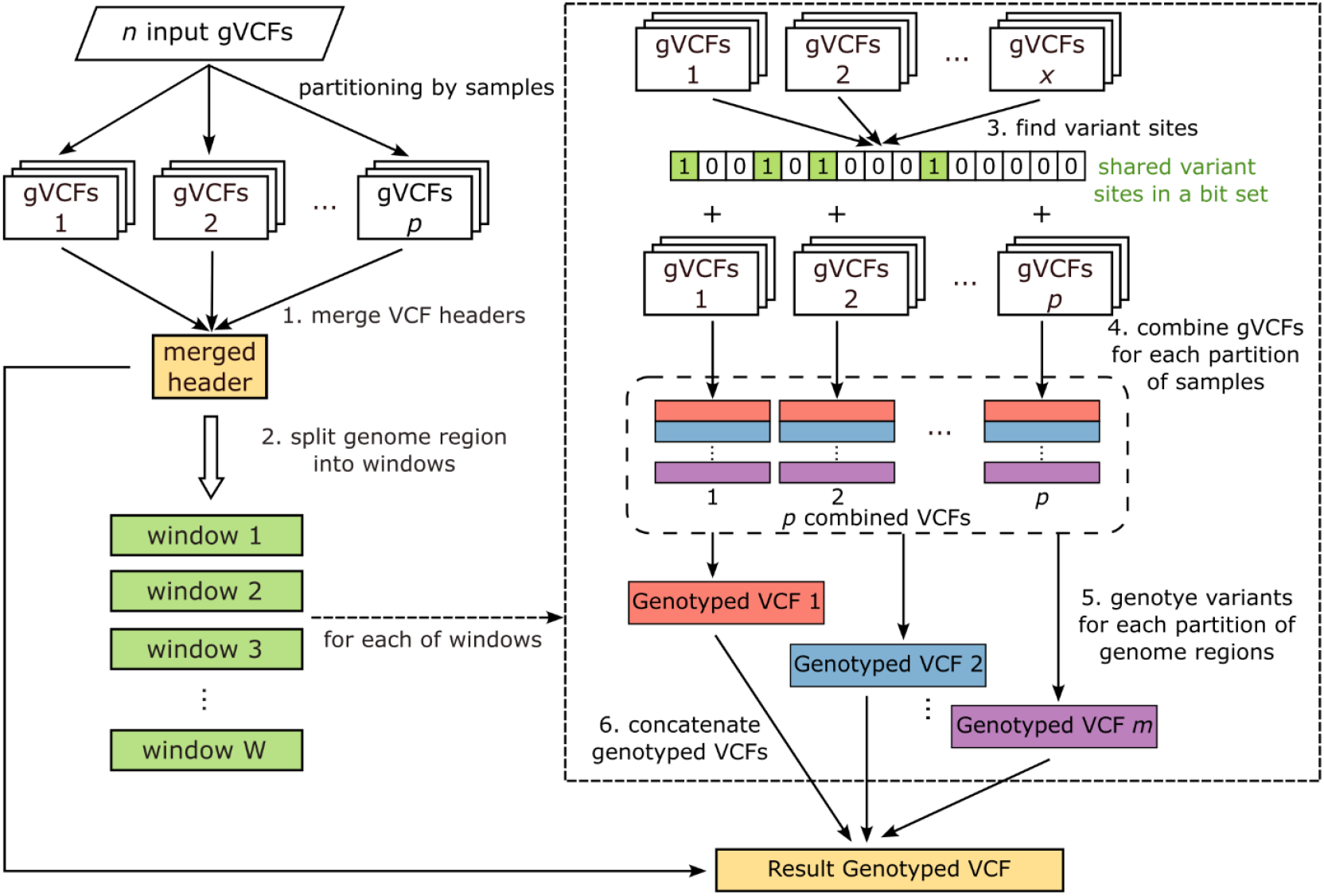
The workflow of DPGT. The input of the method is a list of gVCF files. The method comprises six primary steps, with DPGT parallelizing steps 1 to 5 through the MapReduce technique. In Step 1, DPGT merges gVCF headers into a single header containing all samples from the input gVCF files. DPGT maps the VCF header combiner to partitions of samples (1, 2, …, *p* partitions, where 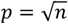 rounded down, each partition has about p samples) to obtain a merged VCF header for each partition and combines these headers into a single VCF header through the reducer. In Step 2, DPGT splits the genome region into windows of a specific size (default: 300Mbp, which means splitting the genome region by chromosome for the human genome). For each window, DPGT performs joint variant calling and outputs genotyped variants to the result VCF file in steps 3 to 6. In Step 3, DPGT identifies variant sites for partitions of samples (1, 2, …, *x* partitions, where *x* equals to the number of virtual cores, each partition has about *n*/*x* samples), storing these sites in bit sets. DPGT obtains shared variant sites (variant sites are indicated in green) by performing logic OR operations on these bit sets. In Step 4, DPGT combines gVCFs for each partition of samples on shared variant sites, resulting in *p* combined VCF files. In Step 5, DPGT genotypes variants by partitioning windows into *m* smaller regions (marked with red, blue and purple in the schematic diagram), and each genotyping worker processes one small region of the *p* combined VCFs. In Step 6, DPGT concatenates the genotyped VCFs to get the result VCF file.

The first key feature contributes to the efficiency of DPGT is its high-concurrency computing workflow. Central to this is the identification of shared variant sites, which are variant sites present in at least one of the input samples (Methods). The input gVCF files are partitioned by sample, with each variant combining task works on one chunk of gVCF files on the same shared variant sites. This variant combining process is highly concurrent, as each combining task runs independently without the need for communication or synchronization between tasks. Because the variant combining tasks are performed on the same shared variant sites, the number of variant records of the combined VCF files (Figure 1. step 4) are identical, and the genomic positions of these variants are consistent. DPGT leverages this feature to perform a second phase of variant combining task efficiently by simply reading variant records from the lines with the same rank in the combined VCFs from the first phase and merging them. The second phase of variant combining is integrated into the genotyper to avoid additional I/O operations involved in writing and reading VCF files. This sample-based task partitioning achieves low memory consumption by eliminating the need to open all gVCF files and load their INFO fields simultaneously in one worker thread. Furthermore, genotyping process was facilitated by executing the tasks in parallel on partitions of genome regions. The genomic region is segmented such that each sub-genomic region contains an equal number of variant sites, a distribution that also depends on the shared variant sites. This segmentation strategy can improve the load balance of genotyping workers, thereby decreasing the overall compute time.

Another key feature that improves the efficiency of DPGT is applying an hybrid method for estimating maximum likelihood expectation (MLE) of the allele counts (MLEAC) and allele frequency (MLEAF). The computing time for MLEAC/MLEAF using GATK best first search algorithm[16] increases linearly with allele counts (Supplementary Figure S1 A) and this makes it a bottleneck for genotyping variants of a large cohort of samples. To speed up this process, DPGT implements a hybrid method, which combines expectation–maximization (EM) algorithm[17] and best first search algorithm, for computing MLEAC/MLEAF(Methods). Benchmarking on chr20:1Mb-5Mb of 2,510 samples from the 1KGP and GIAB datasets demonstrates that the hybrid method, due to 99.79% of EM-steps convergent in less than 20 iterations (Supplementary Figure S1C), reduces computation time by half compared to the GATK best-first search algorithm (Supplementary Figure S1B). Importantly, the hybrid method’s runtime is independent with allele counts (Supplementary Figure S1A), making it highly scalable for large cohorts (Supplementary Figure S1)

#### Computational cost

To evaluate computational cost and scalability on samples of DPGT, GLnexus and GATK (GenomicsDBImport and GenotypeGVCFs tools), joint calling was performed on chromosome 20 of 2,510 1KGP and GIAB samples on the same server with two Intel(R) Xeon(R) Gold 6140 CPU, 187GB of memory and a local SSD. The evaluation was performed on *n*∈{100,1,500,2,510}nested subsets of the 2,510 1KGP and GIAB samples. Although DPGT and GLnexus both natively support multithreading, the two GATK tools are effectively single-threaded. To run GATK on a local computer concurrently, we built a parallelization GATK joint calling workflow based on the developers’ guide using Python multiprocessing pool. Specifically, the chromosome 20 was divided into 65 regions of size 1 Mbp, with each GATK joint calling process handling one region. We ran DPGT and GLnexus using 32 threads and reserving 40GB memory, ran GATK using 20 processes and reserving 160GB memory. DPGT is the fastest among the three evaluated tools. The CPU time of DPGT (96.38 CPU hour) is 26% less than that of GLnexus (130.04 CPU hour) and 81% less than that of GATK (503.96 CPU hour) (n=2,510), with superior scalability to larger cohorts (Figure 2A). We observed similar results using the elapsed time as the evaluation metric (Supplementary Figure S2).

**Figure 2.**
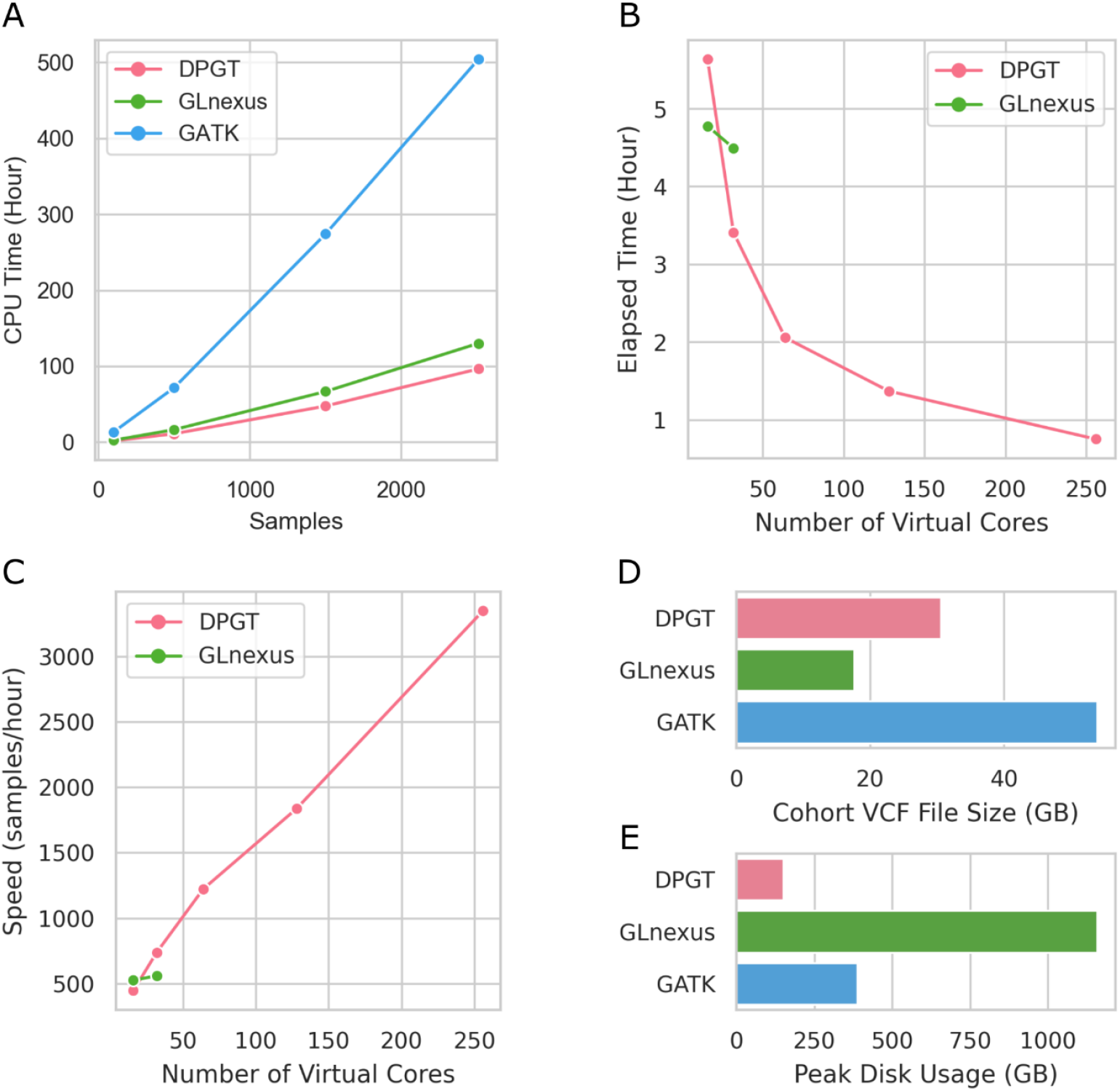
Computational performance evaluation of DPGT. **A** CPU times to combine the chr20 gVCF files into a cohort VCF for *n* ∈ {100, 500, 1000, 1500, 2000, 2510} nested subsets of the 2510 1KGP + GIAB samples, using DPGT, GLnexus and GATK. **B**,**C** Elapsed time (B) and speed (C) for combining the chr20 gVCF files of the 2510 1KGP + GIAB samples, using DPGT and GLnexus with 16, 32, 64, 128 and 256 virtual cores. **D** Cohort VCF file size of DPGT, GLnexus and GATK for chr20 of the 2510 1KGP + GIAB samples. **E** Peak disk usages while running DPGT, GLnexus and GATK on chr20 of the 2510 1KGP + GIAB samples.

To evaluate scalability on multi-cores, we ran DPGT and GLnexus on chromosome 20 of 2,510 1KGP and GIAB samples using 16, 32 virtual cores on the same server and ran DPGT using 64, 128 and 256 virtual cores on a YARN cluster. We did not run GATK in this test, because it does not support multi-threading internally and its runtime is too long (Supplementary Figure S2). Our results show that DPGT has superior scalability to larger number of virtual cores than GLnexus (Figure 2BC). Although DPGT is less efficient than GLnexus using 16 virtual cores, it uses 23% less time elapsed (3.45h) than GLnexus (4.49h) using 32 virtual cores. Moreover, DPGT finishes the job in 45 minutes using 256 virtual cores of a YARN cluster (Figure 2B). The speed of DPGT increases almost linearly with the number of virtual cores (Figure 2C).

To further evaluate the efficiency of DPGT in real-world scenarios, we compared its computational cost with that of GLnexus and GATK on the whole genome of 2,510 1KGP and GIAB samples. DPGT was executed on a YARN cluster using 256 executors (virtual cores), with each executor reserving 7GB of memory. GLnexus was executed on a Sun Grid Engine (SGE) computing cluster, each job analyzed variants on one chromosome. 36 threads and 100GB memory were reserved for each job. Because joint calling on the whole genome of 2,510 samples with GATK is excessively time-consuming, we executed GATK on 1,000 randomly selected genome regions of 100Kbp each, with a total length of 100Mbp. The CPU time for whole genome was estimated by multiplying its CPU time on the randomly selected genome regions by 30. Our results indicate that DPGT offered faster performance than the other two tools, utilizing 33% less effective CPU time than GLnexus and 73% less effective CPU time than GATK (see Table 1). The real elapsed time of DPGT is 45% less than estimated elapsed time of GLnexus and 93% less than estimated elapsed time of GATK (Table 1).

**Table 1.** Computational costs of joint callers on whole genome of 2510 1KGP and GIAB samples.

|  |  | DPGT | GLnexus | GATK (estimated) |
| --- | --- | --- | --- | --- |
| Extract VCF | Effect CPU Time (h) | 0 | 2423.98<sup>a</sup> | 0 |
|  | Reserved CPU Time (h) | 0 | 2706.22 | 0 |
| Joint Calling | Effect CPU Time (h) | 6963.20<sup>b</sup> | 7921.61 | 25621.45 |
|  | Reserved CPU Time (h) | 6963.20 | 9983.95 | 100036.08 |
| Total | Effect CPU Time (h) | 6963.20 | 10345.59 | 25621.45 |
|  | Reserved CPU Time (h) | 6963.20 | 12690.16 | 100036.08 |
| Elapsed Time (h) |  | 27.20 | 49.57<sup>c</sup> | 390.77<sup>c</sup> |

a. GLnexus not supports random fetching variants of target region from VCF files with .tbi index provided, so we extracted variants from target chromosome using bcftools and output VCF files, and then ran GLnexus on them.
b. Because we failed to get effect CPU time from the YARN cluster, we used reserved CPU time as effective CPU time for DPGT.
c. Assuming that GLnexus' and GATK's can operate simultaneously on 256 virtual cores and their CPU usage percents remains unchanged, the elapsed time can be estimated by dividing the reserved CPU time by 256.

We also compared computational cost of DPGT and GLnexus using 9,158 internal whole genome sequencing (WGS) samples and 7 GIAB samples (9,165 samples in total). Joint calling on this large WGS cohort, DPGT used 31% less effective CPU time than GLnexus and the real elapsed time of DPGT(83.5h) is 37% less than estimated elapsed time of GLnexus(132.77h) (Supplementary Table S2). To further evaluate the feasibility of using DPGT for joint calling on larger cohorts, we conducted simulated experiments. We simulated variants (in GRCh38 chr1:10,000,000-11,000,000, which covers 1Mbp genome positions) for 100,000 samples based on allele frequencies derived from gnomAD v3.1.2[18] WGS, on the assumption that each variant is independent and are in Hardy-Weinberg equilibrium (Methods). We ran DPGT on this simulated large cohort using 160 executors, each executor reserved 10GB memory, on a YARN cluster. DPGT finishes this job in 2.65 hour.

One additional cost for running bioinformatics pipelines is storage usage. In our study, the size of the block-compressed gzip file for the final population VCF of chromosome 20 for 2,510 1KGP and GIAB samples generated by DPGT is 30.58GB, which is 43% smaller than the file size generated by GATK (54GB), but 73% larger than the file size generated by GLnexus (17.66GB) (Figure 2C). The difference in output VCF file size between DPGT and GATK is due to their different compression levels. DPGT used compression level 5, while GATK defaults to using compression level 2. When we recompressed GATK’s output file using compression level 5, the resulting VCF file size is 30.88GB, which is very close to the size of the file generated by DPGT. The final cohort VCF file size generated by DPGT and GATK is larger than that generated by GLnexus because they calculate and output all the annotations in INFO field that are useful for downstream analyses, such as Variant Quality Score Recalibration (VQSR) and hard filtering, whereas GLnexus only outputs allele frequency (AF) and allele quality (AQ). However, during joint calling, the peak disk usage for DPGT is much smaller than that for GLnexus and GATK. The peak disk usage for running DPGT (150.43GB) is only 13% and 39% that of running GLnexus (1159.28GB) and GATK (387.87GB) (Figure 2D), respectively.

#### Accuracy

We used several measures to compare the accuracy of DPGT to other joint callers, including comparison to 7 GIAB curated datasets, mendelian error rate in trios, false discovery rate estimated by transmission of alleles in trios, ratio of transitions to transversions (Ti:Tv ratio), and fraction of variants with low HWE p-value. The evaluation was performed by running the three joint callers on 1,000 random selected genome regions of 2,510 1KGP and GIAB samples. Each of the genome regions is 100Kbp long, which resulted in a total length of 100 Mbp.

We assessed the accuracy of raw output generated by three joint variant callers. Our results show that DPGT calls more variants than GLnexus, but less variants than GATK (Table 2). However, DPGT yields the highest recall for both SNP and INDEL variants across GIAB curated datasets (Table 2). The expected ratio of SNP transitions to transversion is roughly 2.0-2.1 in human’s genome wide. Raw call sets have a lower Ti:Tv ratio, while the DPGT call set is much closer to 2.0–2.1 (Table 2). GLnexus’ raw call set has less genotype errors than other caller, it has lower low HWE p value ratio in the population, lower mendelian error rate and false discovery rate (FDR) in trios (Table 2). Our results also shows GATK’s relatively lower INDEL accuracy, as indicated by lower recall and precision values in the GIAB curated datasets compared to the two other joint callers.

**Table 2.**
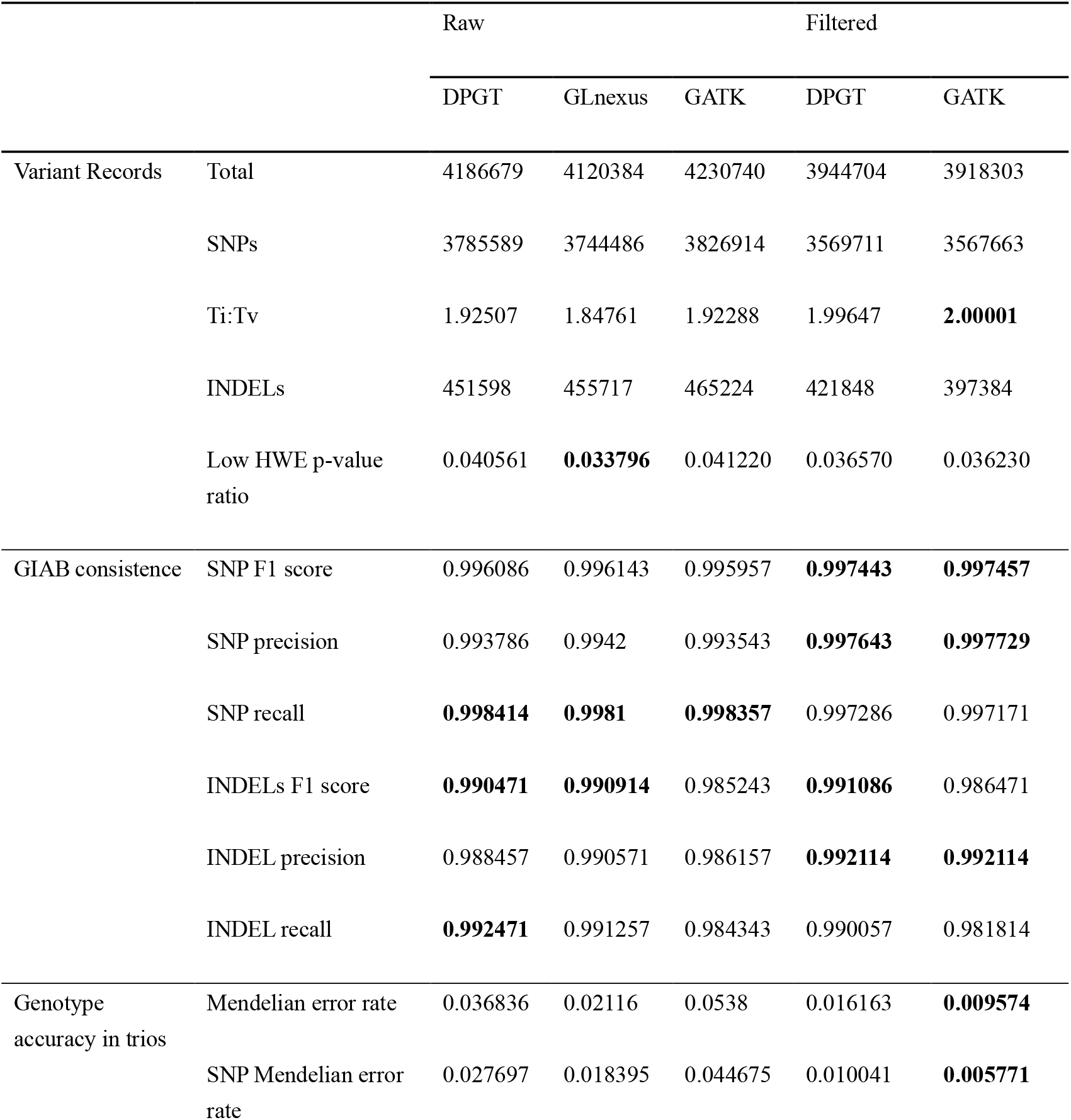
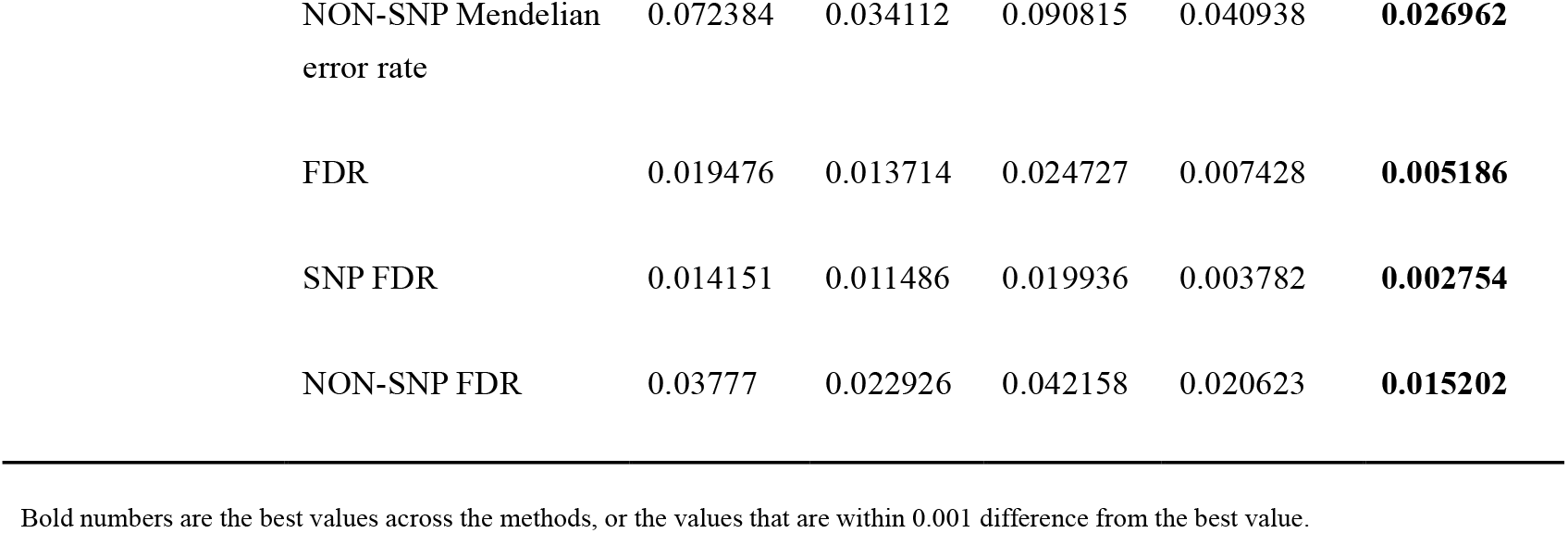
Comparison of accuracy of DPGT and other two joint callers on 1000 random selected regions.

DPGT generates annotations that are comparable to those produced by GATK, enabling us to improve the accuracy of DPGT’s call set. In contrast, GLnexus’s raw result had been filtered internally using specific thresholds such as min_AQ1, min_AQ2, min_GQ, min_allele_copy_number, and min_assumed_allele_frequency [14], and thus, we did not apply any additional filtering to its call set. As an example of using annotations in INFO field, we performed hard filtering on the raw call sets of DPGT and GATK based on GATK’s documentation. Our results shows that both DPGT and GATK filtered call sets have a Ti:Tv ratio that are very close to 2.0-2.1 (Table 2). Hard filtering results in a reduction of 5.7% of SNPs in DPGT call set and 6.8% of SNPs in GATK call set. Nonetheless, the recall in GIAB curated datasets only decreases by 0.11% for DPGT and by 0.12% for GATK. Furthermore, hard filtering improves the precision and F1 scores of both DPGT and GATK for GIAB curated datasets. Additionally, hard filtering decreases the low HWE p-value ratio in the population, mendelian error rate, and false discovery rate (FDR) in trios. Compared to the GIAB curated datasets, DPGT’s filtered call set had similar SNP accuracy to that of GATK’s filtered call set, with better INDEL accuracy.

We also evaluated the accuracy of DPGT and GLnexus on whole genome of 2,510 1KGP and GIAB samples by comparing their raw call sets to GIAB curated datasets. Consistent with the results in randomly selected regions, DPGT has a higher recall than GLnexus, but a lower precision than GLnexus, and their F1 scores are comparable (see Table 3).

**Table 3.** Comparison of accuracy of DPGT and GLnexus on whole genome.

| Method | SNP F1 | SNP Recall | SNP Precision | INDEL F1 | INDEL Recall | INDEL Precision |
| --- | --- | --- | --- | --- | --- | --- |
| DPGT | **0.995357** | **0.998257** | 0.992486 | 0.989586 | **0.992700** | 0.986500 |
| GLnexus | 0.995350 | 0.997967 | **0.992750** | **0.990150** | 0.991750 | **0.988550** |

## Discussion

In order to meet the demand for variant joint calling in large-scale population studies, we implemented a novel distributed joint calling framework within the DPGT software. DPGT significantly outperforms GATK in speed. It is also faster than GLnexus, even while computing and outputting more valuable annotation information. It offers excellent sample and multi-core scalability. DPGT simplifies joint calling tasks for large cohorts with a single command, eliminating the need for users to create complex parallel workflows. Furthermore, its accuracy is on par with other software solutions. The efficiency of DPGT is attributed to its high-concurrency computing framework and the implementation of a hybrid method for calculating MLEAC/MLEAF. Using shared variant sites DPGT makes it possible to combine variants from partition of samples independently and improves the load balance. The hybrid MLEAC/MLEAF computing method improves the efficiency and the scalability to large cohorts.

DPGT’s computing workflow has three distinct advantages over other approaches. First, it divides the joint calling task by two dimensions, the sample dimension and the genome position dimension. This dual-dimensional division improves the utilization of multiple cores within a computing cluster, a feature not presents in GATK or GLnexus (which only divides the joint calling task by samples, also known as cohort-sharding). Second, DPGT employs VCF file streams when performing variant combination, as opposed to a disk-based database. This approach reduced I/O activity compared to GATK’s GenomicsDB and GLnexus’s RocksDB. Lastly, DPGT’s memory usage is low due to its task division strategy for parallel processing and the implementation of low memory footprint data structures, such as bit-set.

We have shown that DPGT already represents an efficient tool for joint calling of large cohorts. However, we think that further improvements are still possible. DPGT only supports inputs from GATK HaplotypeCaller, Sentieon and Illumina DRAGEN, additional efforts are required to support inputs from other state-of-art variant callers like DeepVariant and Streka2. Furthermore, recent studies have shown that pan-genome graphs can improve the accuracy of sequence alignment and variant calling[19–21]. Using DPGT in conjunction with pan-genome graph-based variant calling methods may improve the accuracy.

As NGS gets more economically feasible, large-scale population genetics study could be more and more common in the near future. Data analysis occupies a substantial portion of the time and economic costs in such studies. In order to reduce these costs, we provide an open-source software for fast, scalable, and accurate joint variant calling.

## Methods

### Finding variant sites

One of the key innovations of DPGT is the pre-computation of shared variant sites. Shared variant site is defined as a genome position where there is at least one true alternative allele (the ALT field in VCF file is not null (.) or <NON_REF>). SNP and insertion variants cover one variant site. A deletion variant covers multiple contiguous variant sites, the covered length is equal to the deletion size. The variant sites from all input gVCFs are shared among all combining tasks and are thus referred to as shared variant sites. DPGT uses bit set structure to store variant sites, where the index of the bit set represents the relative position in genome and the value (0 or 1) of the bit set represents if the position is a variant site. The compact memory footprint of the bit set structure, combined with efficient merging through logical OR operations, enables effective handling of variant sites. For the longest chromosome of GRCh38 (chr1 248956422 bp), the shared variant sites bit set consumes 29.67 MB memory.

### Combining Variants

DPGT implements a two phases method to combining variants of input gVCF files. At the first phase, DPGT combines variants for partitions of samples (Figure 1, step 4), which outputs *p* (the number of partitions) combined VCF files. At the second phase, DPGT combines variants of the *p* combined VCF files to get combined variants for all of the input samples. The core algorithm for merging variants[6] is the same for the two phase, but the algorithms for grouping variants of a genome position are different. For the first phase, DPGT implements a plane sweep algorithm for combining variants on shared variant sites. It read variants of the *q* input gVCFs in genome position order, put the variant into a linked-list if this variant’s genome interval is overlapping with current variant site, otherwise it combines variants on current variant site by using GATK’s variants merging algorithm[6], then it moves to the next variant site and remove variants of which genome interval not overlapping with the variant site from the linked list. For the second phase, grouping variants of a genome position is simpler and more efficient. Owing to the using of shared variant sites, the number of variant records of the *p* combined VCF files (Figure 1. step 4) are identical, and the genomic positions of these variants are consistent. DPGT simply need to read variant records from the lines with the same rank from the *p* combined VCF files and merge them using the same algorithm as the first phase. Both variant grouping algorithms have a low memory footprint and low I/O (Input/Output, reading from or writing to a disk) activity. The number of variant records storing in the linked list for grouping variants at the first phase is *O(q)*, where q is the number of the input gVCF files. DPGT reads *p* variants from *p* combined VCF files at a time at the second phase. Both variant grouping algorithms read variant records from VCF file streams and not use database on disk (GenomicsDB for GATK and RocksDB for GLnexus) like other joint calling method does. Need to note that DPGT only merge variants on the shared variant sites at the first combining phase, thereby it does not support incremental joint calling.

### Estimating MLEAC/MLEAF

DPGT implements a hybrid approach for estimating the maximum likelihood expectation (MLE) of the allele counts (MLEAC) and the allele frequency (MLEAF). For variants with small allele count DPGT applies GATK’s best first search algorithm, for variants with large allele count it applies expectation-maximization algorithm. First, DPGT applies 100 rounds of EM iterations to estimate an allele frequency *ψ*. Given we know the estimate *ψ^(t)^* at the t-th iteration, the estimate at the (*t* +1)-th iteration is:

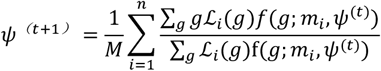

where *ℒ_i_(g)* is compute from PL of i-th sample in the VCF file, *g* is the number of alternative alleles in GT (g = 0 for GT = 0/0, g = 1 for GT = 0/1 and g = 2 for GT = 1/1). *M* = ∑*_i_ m_i_* is the total number of chromosomes in samples. *n* is the number of samples. *f*(*g; m_i_, ψ*) is computed by:

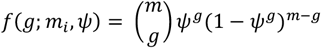

is the probability mass function of the binomial distribution Binom(*m, ψ*). *m* is the ploidy of the sample. The MLEAC is computed by *α = ψM*, rounded up.

Then, if the allele count *α* is less than 100, DPGT switches to GATK’s best first search algorithm. This hybrid approach ensures consistent MLEAC/MLEAF estimation to GATK’s best first search algorithm for smaller allele counts, while significantly expediting the computational process for larger allele counts.

### Simulating variants

To evaluate the practicality of conducting joint calling on large cohorts using DPGT, we simulated variants in GRCh38 chr1:10,000,000-11,000,000 for 100,000 samples. The simulation was based on the population VCF which contains 76,156 whole genomes from gnomAD v3.1.2, with the following steps implemented. Firstly, we used bcftools[22] to extract biallelic variants in the genome region chr1:10,000,000-11,000,000 from gnomAD v3.1.2 VCF. Secondly, assuming the genotypes of each variant to be in Hardy-Weinberg equilibrium, we calculated the probabilities of genotypes 0/0, 0/1, and 1/1 based on the allele frequencies obtained from step one by:

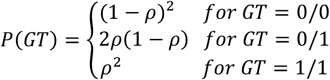

where *p* is the alt allele frequency.

After that, we uniformly randomly selected a genotype from genotypes 0/0, 0/1, and 1/1 based on the calculated probabilities as the simulated genotype for each variant. Thirdly, we generated a consensus sequence by applying the simulated variants to GRCh38 reference sequence of chr1:10,000,000-11,000,000 using bcftools consensus and simulated 30x depth 150bp paired-end reads using art_illumina. Lastly, we generated variants for each sample by mapping the simulated reads to GRCh38 reference using bwa mem and called variants using GATK HaplotypeCaller. We repeated steps two to four 100,000 times using different seeds to generate a gVCF file for each sample.

### Variant quality assessment

The basic variant statistics of the joint calling data sets, including total number of variants, number of SNP variants, number of INDEL variants and ratio of transitions to transversions (Ti:Tv ratio) of SNPs, were computed using bcftools. The Ti:Tv ratio represented here is the average Ti:Tv ratio of all samples from the call set. Deviations from HWE can be indicative of recurrent artifacts in a variant calling algorithm[23]. We used vcftools[24] to calculate the Hardy-Weinberg equilibrium (HWE) p-value[25] of biallelic SNPs and determined the proportion of variants with a p-value less than 10^-5^. We compare joint calling datasets to GIAB truth sets (V3.3.2) using RTG vcfeval[26], the evaluation region is the overlapped regions of joint calling regions and GIAB high-confident regions. Before running RTG vcfeval, we filtered out homozygous reference (0/0) and no called (./.) variants, trimmed not used alternative alleles, and did normalization on alleles using bcftools.

The Mendelian error rate was estimated as the fraction of variants that violates Mendelian low in trios. We assumed that the alleles transmit from parent to child with equal likelihood and used the transmission rate (TMR) to estimate false discovery rate(FDR). More info about the method was described[19]. The trios evaluated here were two trios from GIAB (HG002: child, HG003: father, HG004: mother, all of Ashkenazi Jews ancestry; HG005: child, HG006: father, HG007: mother, all of Chinese ancestry) and one cryptic trio from 1KGP (NA20900: child, NA20891: father, NA20882: mother, all of Gujarati Indian ancestry).

## Availability of source code and requirements

- Project name: DPGT
- Project home page: https://github.com/BGI-flexlab/DPGT
- Operating system(s): Linux
- Programming language: Java, C++
- Other requirements: Boost C++ library 1.74.0 or higher, Java 1.8, Spark 2.4.0 or higher.
- License: GPLv3
- Any restrictions to use by non-academics: No Restriction

## Data availability

For 2504 1KGP samples, high-coverage sequencing reads in cram format is available at 1KGP FTP website (cram file URL for each of samples is list in [27]), and the mean depth of the samples is about 30x. The pipeline to generate these cram files is well documented in the pipeline description[28].

Sample NA12878(HG001) is included in 2504 1KGP samples, other 6 GIAB samples (HG002-HG007) high-coverage sequencing reads produced by MGI platform are available at GIAB FTP website[29](detailed in Supplementary Table S4). The seven GIAB V3.3.2 truth sets VCF and high confident region bed files are available at GIAB FTP website[30].

## Abbreviations

1KGP: 1000 Genome Project
AC: allele count
AF: allele frequency
AQ: qualities for alternate alleles
BAM: binary sequence alignment and mapping format
EM: expectation–maximization
FDR: false discovery rate
gVCF: genome variant calling format
GIAB: Genome In the Bottle Project
HWE: Hardy-Weinberg equilibrium
INDEL: small insertion and deletions
MLEAC: maximum likelihood expectation allele count
MLEAF: maximum likelihood expectation allele frequency
NON-REF: non-reference alternate allele, this allele used as a wild-card to match any alternate allele
RCM: reference confidence model
SNP: single nucleotide polymorphism
TMR: transmission rate
VCF: variant calling format
WGS: whole genome sequencing

## Authors’ contributions

Yuxiang Li supervised the research. Yong Zhang and Chun Gong conceptualized the software. Chun Gong, Qi Yang, and Ruiwen Wan implemented the software code. Qi Yang and Shengkang Li analyzed the data. Chun Gong, Qi Yang, and Shengkang Li contributed to the writing of the manuscript. All the authors read and approved the final manuscript.

## Acknowledgements

We would like to thank DCS Cloud (https://cloud.stomics.tech/) for providing the computational resources and software support necessary for this study.

