## Supplementary Material for "DPGT: A spark based high-performance joint variant calling tool for large cohort sequencing"

#### 1. Supplementary Figures

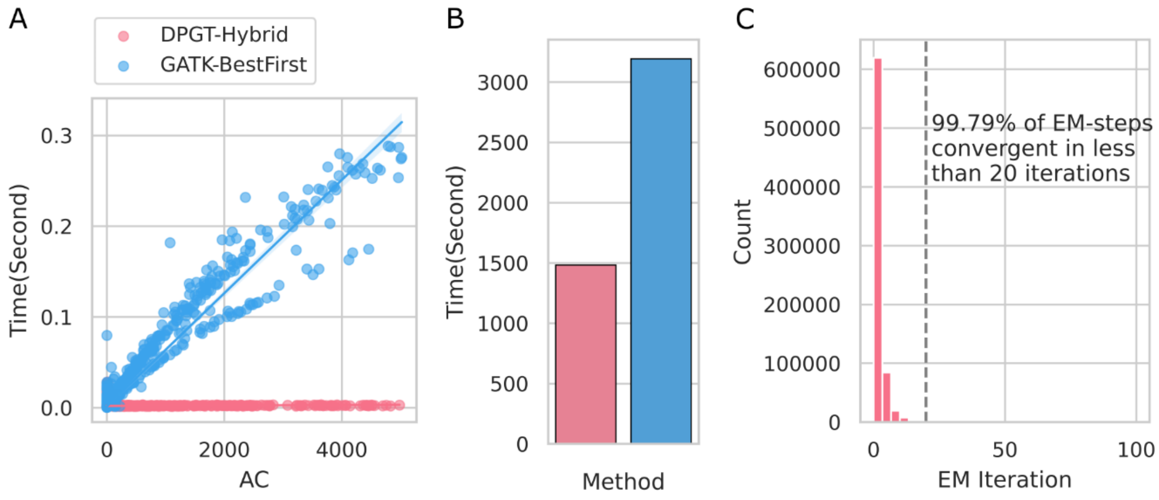


**Supplementary Figure S1.** Comparison of GATK’s best first search algorithm and DPGT’s hybrid algorithms for calculating MLEAC/MLEAF. Joint calling was run on chr20:1Mb-5Mb of 2,510 1KGP and GIAB samples using the two methods, the allele counts, EM iterations and time for compute MLEAC/MLEAF were recorded for analysis. **A** Correlation of time for calculating MLEAC/MLEAF and allele count (AC). **B** Total time for calculating MLEAC/MLEAF of the two methods. **C** The number of iterations required for the Expectation-Maximization (EM) algorithm to achieve convergence.

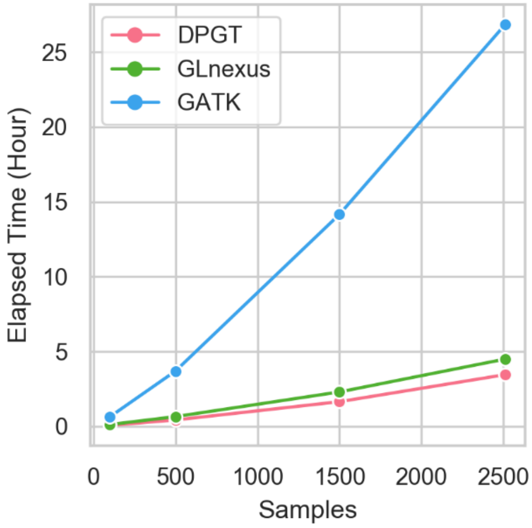


**Supplementary Figure S2.** Elapsed time to combine the chr20 gVCF files into a cohort VCF for $n \in \{100, 500, 1{,}000, 1{,}500, 2{,}000, 2{,}510\}$ nested subsets of the 2, 510 1KGP + GIAB samples, using DPGT, GLnexus and GATK.

#### 2. Supplementary Tables

**Supplementary Table S1. Comparison of joint calling tool features.**

| Feature | DPGT | GLnexus | GATK |
|---|---|---|---|
| Freely available | Yes | Yes | Yes |
| Local computer | Yes | Yes | Yes |
| Computer cluster | Yes | No[a] | No |
| Multithreading | Yes | Yes | No[b] |
| Resume job after interrupt | Yes | No | No |
| Random analysis target region of genome | Yes | No | Yes |
| Final cohort VCF compatible with GATK hard-filter and VQSR | Yes | No | Yes |
| Support input gVCF from DeepVariant, xAtlas, weCall | No | Yes | No |

a. The open source GLnexus lack some features for production usage and cannot be run on a compute cluster without split input files and jobs by genome region.

b. While GATK GenomicsDBImport and GenotypeGVCFs do use multiple threads for reading or writing VCF files, their core functionalities are single threaded.

**Supplementary Table S2. Comparison of DPGT and GLnexus on 9,158 internal and 7 GIAB samples**

| Software | | DPGT | GLnexus |
|---|---|---|---|
| Extract VCF | Effect CPU Time (h) | 0 | 5,477.319[a] |
| | Reserved CPU Time (h) | 0 | 5,614.063 |
| Joint Calling | Effect CPU Time (h) | 21,376[b] | 25,577.06 |
| | Reserved CPU Time (h) | 21,376 | 28,375.51 |
| Total | Effect CPU Time (h) | 21,376 | 31,054.38 |
| | Reserved CPU Time (h) | 21,376 | 33,989.57 |
| | Elapsed Time (h) | 83.5 | 132.77[c] |

a. GLnexus does not supports random fetching variants of target region from VCF files with .tbi index provided, so we extracted variants from target chromosome using bcftools and output VCF files, and then ran GLnexus on them.

b. Because we failed to get effect CPU time from the YARN cluster, we used reserved CPU time as effective CPU time for DPGT.

c. GLnexus' elapsed time were estimated by dividing their reserved CPU time by 256.

**Supplementary Table S3. Hardware resources of the YARN cluster that used for running DPGT.**

| Node | Architecture | CPU (s) | Core(s) per socket | Socket (s) | Model name | Memory (physical) | Memory (Yarn) | Virtual Cores (Yarn) |
|---|---|---|---|---|---|---|---|---|
| 1 | x86_64 | 48 | 12 | 2 | Intel(R) Xeon(R) CPU E5-2650 v4 @ 2.20GHz | 251G | 210G | 38 |
| 2 | x86_64 | 48 | 12 | 2 | Intel(R) Xeon(R) CPU E5-2650 v4 @ 2.20GHz | 251G | 210G | 38 |
| 3 | x86_64 | 48 | 12 | 2 | Intel(R) Xeon(R) CPU E5-2650 v4 @ 2.20GHz | 251G | 210G | 38 |
| 4 | x86_64 | 48 | 12 | 2 | Intel(R) Xeon(R) CPU E5-2650 v4 @ 2.20GHz | 251G | 210G | 40 |
| 5 | x86_64 | 48 | 12 | 2 | Intel(R) Xeon(R) CPU E5-2650 v4 @ 2.20GHz | 251G | 210G | 38 |
| 6 | x86_64 | 32 | 8 | 2 | Intel(R) Xeon(R) CPU E5-2620 v4 @ 2.10GHz | 125G | 110G | 24 |
| 7 | x86_64 | 48 | 12 | 2 | Intel(R) Xeon(R) CPU E5-2650 v4 @ 2.20GHz | 251G | 210G | 38 |
| 8 | x86_64 | 48 | 12 | 2 | Intel(R) Xeon(R) CPU E5-2650 v4 @ 2.20GHz | 251G | 210G | 40 |
| 9 | x86_64 | 48 | 12 | 2 | Intel(R) Xeon(R) CPU E5-2650 v4 @ 2.20GHz | 251G | 190G | 38 |
| 10 | x86_64 | 48 | 12 | 2 | Intel(R) Xeon(R) CPU E5-2650 v4 @ 2.20GHz | 251G | 210G | 38 |
| 11 | x86_64 | 48 | 12 | 2 | Intel(R) Xeon(R) CPU E5-2650 v4 @ 2.20GHz | 251G | 210G | 38 |
| 12 | x86_64 | 48 | 12 | 2 | Intel(R) Xeon(R) CPU E5-2650 v4 @ 2.20GHz | 251G | 210G | 38 |
| 13 | x86_64 | 32 | 8 | 2 | Intel(R) Xeon(R) CPU E5-2620 v4 @ 2.10GHz | 125G | 110G | 24 |
| 14 | x86_64 | 48 | 12 | 2 | Intel(R) Xeon(R) CPU E5-2650 v4 @ 2.20GHz | 251G | 210G | 38 |

**Supplementary Table S4. GIAB HG002-HG007 WGS sequences data URLs.**

| Sample | READ1 URL | READ2 URL |
|---|---|---|
| HG002 | https://ftp-trace.ncbi.nlm.nih.gov/ReferenceSamples/giab/data/AshkenazimTrio/HG002_NA24385_son/MGISEQ/PCR-free/NA24385/MGISEQ2000_PCR-free_NA24385_V100002807_L03_1.fq.gz | https://ftp-trace.ncbi.nlm.nih.gov/ReferenceSamples/giab/data/AshkenazimTrio/HG002_NA24385_son/MGISEQ/PCR-free/NA24385/MGISEQ2000_PCR-free_NA24385_V100002807_L03_2.fq.gz |
| HG003 | https://ftp-trace.ncbi.nlm.nih.gov/ReferenceSamples/giab/data/AshkenazimTrio/HG003_NA24149_father/MGISEQ/PCR-free/NA24149/MGISEQ2000_PCR-free_NA24149_V100002807_L01_1.fq.gz | https://ftp-trace.ncbi.nlm.nih.gov/ReferenceSamples/giab/data/AshkenazimTrio/HG003_NA24149_father/MGISEQ/PCR-free/NA24149/MGISEQ2000_PCR-free_NA24149_V100002807_L01_2.fq.gz |
| HG004 | https://ftp-trace.ncbi.nlm.nih.gov/ReferenceSamples/giab/data/AshkenazimTrio/HG004_NA24143_mother/MGISEQ/PCR-free/NA24143_1/MGISEQ2000_PCR-free_NA24143_1_V100003043_L03_1.fq.gz | https://ftp-trace.ncbi.nlm.nih.gov/ReferenceSamples/giab/data/AshkenazimTrio/HG004_NA24143_mother/MGISEQ/PCR-free/NA24143_1/MGISEQ2000_PCR-free_NA24143_1_V100003043_L03_2.fq.gz |
| HG005 | https://ftp-trace.ncbi.nlm.nih.gov/ReferenceSamples/giab/data/ChineseTrio/HG005_NA24631_son/MGISEQ/PCR-free/NA24631/MGISEQ2000_PCR-free_NA24631_V100002812_L01_1.fq.gz | https://ftp-trace.ncbi.nlm.nih.gov/ReferenceSamples/giab/data/ChineseTrio/HG005_NA24631_son/MGISEQ/PCR-free/NA24631/MGISEQ2000_PCR-free_NA24631_V100002812_L01_2.fq.gz |
| HG006 | https://ftp-trace.ncbi.nlm.nih.gov/ReferenceSamples/giab/data/ChineseTrio/HG006_NA24694-huCA017E_father/BGISEQ500/BGISEQ500_PCRfree_NA24694_CL100076304_L01_read_1.fq.gz | https://ftp-trace.ncbi.nlm.nih.gov/ReferenceSamples/giab/data/ChineseTrio/HG006_NA24694-huCA017E_father/BGISEQ500/BGISEQ500_PCRfree_NA24694_CL100076304_L01_read_2.fq.gz |
| HG007 | https://ftp-trace.ncbi.nlm.nih.gov/ReferenceSamples/giab/data/ChineseTrio/HG007_NA24695-hu38168_mother/BGISEQ500/BGISEQ500_PCRfree_NA24695_CL100076245_L01_read_1.fq.gz | https://ftp-trace.ncbi.nlm.nih.gov/ReferenceSamples/giab/data/ChineseTrio/HG007_NA24695-hu38168_mother/BGISEQ500/BGISEQ500_PCRfree_NA24695_CL100076245_L01_read_2.fq.gz |

### 3. Supplementary Notes

**Supplementary Note 3.1. SNP and INDEL calling for individual of 1KGP**

We used sentieon HaplotypeCaller (version: sentieon-genomics-201911) for SNP and INDELL calling of individual by inputting cram files. The command we ran was the following:

```
sentieon driver -r hg38.fa \
    -t 8 \
    -i sample.cram \
    --algo Haplotyper \
    -d dbsnp_138.hg38.vcf.gz \
    --emit_mode gvcf \
    sample.g.vcf.gz
```

**Supplementary Note 3.2. SNP and INDEL calling for individual of GIAB**

We ran sentieon (version: sentieon-genomics-202010.01) bwa mem and HaplotypeCaller pipeline to call SNP and INDELL of GIAB samples (HG002 to HG007) following sentieon's documentation (https://support.sentieon.com/versions/202010.01/manual/DNAseq_usage/dnaseq/). We skipped Indel realignment step because it is optional. We also skipped Base quality score recalibration (BQSR) step because the model did not support MGI platform. The command we ran was the following:

```
sentieon bwa mem -R "@RG\tID:${SAMPLE}\tSM:${SAMPLE}\tPL:MGI" \
    -t 16 hg38.fa sample.R1.fastq.gz sample.R2.fastq.gz \
   | sentieon util sort -r hg38.fa -o sample.sorted.bam \
   -t 16 --sam2bam -i -

sentieon driver -t 16 -i sample.sorted.bam \
    --algo LocusCollector --fun score_info sample.markdup.score

sentieon driver -t 16 -i sample.sorted.bam \
    --algo Dedup --score_info sample.markdup.score \
    --metrics sample.dedup.metrics sample.dedup.bam

sentieon driver -t 16 -r hg38.fa -i sample.dedup.bam --algo
Haplotyper \
    -d dbsnp_138.hg38.vcf.gz --emit_mode gvcf sample.g.vcf.gz
```

**Supplementary Note 3.3. Joint calling using DPGT on local computer**

We used `--master local[thread]` option of `spark-submit` to run DPGT on local computer, where `thread` is the number of threads to be use. DPGT is implemented using C++ and Java, it requires the compiled C++ dynamic library to be in `LD_LIBRARY_PATH`. jemalloc was used for reducing memory fragmentation.

```
export  LD_LIBRARY_PATH=/path_to/dpgt_dir/build/lib:${LD_LIBRARY_PATH}
export LD_PRELOAD=/path_to/jemalloc_5.3.0/lib/libjemalloc.so
spark-submit \
    --conf spark.dynamicAllocation.enabled=false \
    --conf spark.memory.fraction=0.01 \
    --conf spark.memory.storageFraction=0.01 \
    --master local[${THREAD}] \
    --driver-memory 60g \
    --class org.bgi.flexlab.dpgt.jointcalling.JointCallingSpark \
    dpgt.jar \
    -i vcfs.list \
    -r hg38.fasta \
    -o result -n 50 -j ${THREAD} \
    -s 10.0 \
    --max-alternate-alleles 10
```

Where vcfs.list is a text file of paths to gVCF files, one file path per line.

**Supplementary Note 3.4. Joint calling using DPGT on yarn cluster**

We used `--master yarn` option of spark-submit to run DPGT on YARN cluster. The `spark.executor.extraLibraryPath` and `spark.driver.extraLibraryPath` configurations were set to be the value of `LD_LIBRARY_PATH`, `spark.excutorEnv.LD_PRELOAD` was set to be the path of jemalloc library. These configurations enable the Spark driver and executor process to load the DPGT C++ dynamic library and utilize jemalloc.

```
export LD_LIBRARY_PATH=/path_to/dpgt_dir/build/lib:${LD_LIBRARY_PATH}
export LD_PRELOAD=/path_tojemalloc_5.3.0/lib/libjemalloc.so
spark-submit \
    --conf spark.dynamicAllocation.enabled=false \
    --conf spark.excutorEnv.LD_PRELOAD=${LD_PRELOAD} \
    --conf spark.driver.extraLibraryPath=${LD_LIBRARY_PATH} \
    --conf spark.executor.extraLibraryPath=${LD_LIBRARY_PATH} \
    --conf spark.driver.maxResultSize=10g \
    --conf spark.memory.fraction=0.01 \
    --conf spark.memory.storageFraction=0.01 \
    --master yarn \
    --driver-memory 30g --num-executors ${EXECUTORS} \
    --executor-memory 7g --executor-cores 1 \
    --class org.bgi.flexlab.dpgt.jointcalling.JointCallingSpark \
    dpgt.jar \
    -i vcfs.list \
    -r hg38.fasta \
    -o result -n 50 -j ${EXECUTORS} \
    -s 10.0 \
    --max-alternate-alleles 10
```

**Supplementary Note 3.5. Joint calling on randomly selected 1000 genome regions for 2,510 1KGP and GIAB samples.**

Considering the significant demands of computational resources and time required to perform joint calling on the whole genome of 2,510 samples using GATK, we executed GATK joint calling on randomly selected genomic regions. Each of the genome regions was 100Kbp long, which resulted in a total length of 100 Mbp. To ensure fairness in comparing the accuracy of the joint calling datasets, we also ran DPGT and GLnexus on these genomic regions.

We created 1,000 100Kbp genome regions using python script. This process started by splitting whole genome into small genome regions of 100Kbp long and then 1,000 genome regions were chosen randomly.

We performed joint calling using GATK (version: 4.2.6.1) on randomly selected genome regions following its best practice. Firstly, we built a GenomicsDB with GATK GenomicsDBImport. We set the interval padding to 100 to avoid edge effect on INDEL calling. The command we ran was the following:

```
gatk --java-options "-Xmx40g" GenomicsDBImport \
      --reader-threads 4 \
      --genomicsdb-shared-posixfs-optimizations true \
      --batch-size 50 \                                       --
      tmp-dir ${tmpdir} \
      --genomicsdb-workspace-path ${outdir}/gendb \
      -L ${interval} \
      --interval-padding 100 \
      --sample-name-map \
      ${sample_map} \
      --genomicsdb-vcf-buffer-size 65536
```

Where `sample_map` was the tab-delimited text file of sample names and paths to samples. `interval` was one of the 1,000 randomly selected genome regions. The options `–batch-size` was used for reduce memory consumption.

We then used GATK GenotypeGVCFs to perform joint genotyping. The command we ran was the following:

```
gatk --java-options "-Xmx10g" GenotypeGVCFs \
     -R ${REF} \
      -G StandardAnnotation -G AS_StandardAnnotation \
      --allow-old-rms-mapping-quality-annotation-data \
     -O ${outdir}/result.pad.vcf.gz \
      -L ${interval} --interval-padding 100 \
      -V gendb://${outdir}/gendb

gatk --java-options "-Xmx10g" SelectVariants \
     -V ${outdir}/result.pad.vcf.gz \
      -L ${interval} \
     -O ${outdir}/result.vcf.gz
```

Where `REF` was the GRCh38 reference sequences. We used GATK SelectVariants to select variants in the target region.

For each of the 1,000 randomly selected genome regions, we submitted the GATK joint calling commands to a Sun Grid Engine (SGE) cluster.

bcftools concat was used for combining joint called vcfs for all genome regions. The command we ran was the following:

```
bcftools concat -n -f ${VCFS} -O z -o result.vcf.gz
```

Where `VCFS` was the joint called vcfs for all genome regions.

We used linux `time` command to estimate compute resources. Only GATK GenomicsDBImport and GenotypeGVCFs processes were included for computational resources estimation.

We performed joint calling using GLnexus (GLnexus version: v1.4.1-0-g68e25e5-dirty, jemalloc version: 5.3.0) on randomly selected genome regions in two steps. Firstly, we extracted variants of the genome region from 2,510 gVCF files using bcftools view. We used python multiprocess pool to run multiple variants extracting job simultaneously. We used 4 processes to run the variants extracting jobs. The bcftools view command we ran was the following:

```
bcftools view -r ${region} \
     -O z -o sample.region.g.vcf.gz sample.g.vcf.gz
tabix -f sample.region.g.vcf.gz
```

Where `region` was the target genome region.

We then ran GLnexus on gVCFs of the target genome region using following command:

```
glnexus_cli \
    --dir ${outdir}/GLnexus.DB \
    --bed ${outdir}/target.bed --list region.vcfs.list  \
    --threads 4 --mem-gbytes 10 \
    | bcftools view --threads 4 -O z -o ${outdir}/result.vcf.gz
```

Where `target.bed` was the target genome region, `region.vcfs.fofn` was a text file of paths to gVCF files for the target genome region.

For each of the 1,000 randomly selected genome regions, we submitted the GLnexus joint calling commands to a Sun Grid Engine (SGE) cluster.

We performed joint calling using DPGT (version: 1.2.8.0) on randomly selected genome regions. The commands we ran was the following:

```
export  LD_LIBRARY_PATH=/path_to/dpgt_dir/build/lib:${LD_LIBRARY_PATH}
export LD_PRELOAD=/path_to/jemalloc_5.3.0/lib/libjemalloc.so
spark-submit \
    --conf spark.dynamicAllocation.enabled=false \
    --conf spark.memory.fraction=0.01 \
    --conf spark.memory.storageFraction=0.01 \
    --master local[4] \
    --driver-memory 10g \
    --class org.bgi.flexlab.dpgt.jointcalling.JointCallingSpark \
    dpgt.jar \
    -i vcfs.list \
    -r ${REF} \
    -o result -n 50 -j 4 \
    -s 10.0 \
    --max-alternate-alleles 10 \
```

```
    -l ${region}
```

Where REF was the GRCh38 reference sequences, vcfs.list was a text file of paths to gVCFs of 2510 samples, region was the target genome region.

For each of the 1,000 randomly selected genome regions, we submitted the DPGT joint calling command to a Sun Grid Engine (SGE) cluster.

**Supplementary Note 3.6. Joint calling on whole genome of 2,510 1KGP and GIAB samples.**

We performed joint calling using DPGT (version: 1.2.8.0) on whole genome of 2,510 1KGP and GIAB samples using 256 virtual cores of a YARN cluster. The command we ran was the following:

```
export LD_LIBRARY_PATH=/path_to/dpgt_dir/build/lib:${LD_LIBRARY_PATH}
export LD_PRELOAD=/path_tojemalloc_5.3.0/lib/libjemalloc.so
spark-submit \
    --conf spark.dynamicAllocation.enabled=false \
    --conf spark.excutorEnv.LD_PRELOAD=${LD_PRELOAD} \
    --conf spark.driver.extraLibraryPath=${LD_LIBRARY_PATH} \
    --conf spark.executor.extraLibraryPath=${LD_LIBRARY_PATH} \
    --conf spark.driver.maxResultSize=10g \
    --conf spark.memory.fraction=0.01 \
    --conf spark.memory.storageFraction=0.01 \
    --master yarn \
    --driver-memory 30g --num-executors 256 \
    --executor-memory 7g --executor-cores 1 \
    --class org.bgi.flexlab.dpgt.jointcalling.JointCallingSpark \
    dpgt.jar \
    -i vcfs.list \
    -r hg38.fasta \
    -o result -n 50 -j 256 \
    -s 10.0 \
    --max-alternate-alleles 10
```

Where vcfs.list was a text file of paths to gVCFs of 2, 510 samples.

We performed joint calling using GLnexus (GLnexus version: v1.4.1-0-g68e25e5-dirty, jemalloc version: 5.3.0) on whole genome of 2,510 1KGP and GIAB samples by splitting jobs by chromosomes. Firstly, we extracted variants of the target chromosome from 2,510 gVCF files using bcftools view. We used python multiprocess pool to run multiple variants extracting job simultaneously. 36 processes were used to extract the variants. The bcftools view command we ran was the following:

```
bcftools view -r ${chrom} \
      -O z -o sample.region.g.vcf.gz sample.g.vcf.gz
tabix -f sample.region.g.vcf.gz
```

Where `chrom` was the target chromosome.

We then ran GLnexus on gVCFs of the target chromosome using following command:

```
glnexus_cli \
    --dir ${outdir}/GLnexus.DB \
    --bed ${outdir}/target.bed --list chrom.vcfs.list  \
    --threads 36 --mem-gbytes 100 \
    | bcftools view --threads 12 -O z -o ${outdir}/result.vcf.gz
```

Where `target.bed` was the target chromosome region, `chrom.vcfs.list` was a text file of paths to gVCF files for the target chromosome region.

For each of the 25 chromosomes (chr1, chr2, …, chrX, chrY, chrM), we submitted the GLnexus joint calling commands to a Sun Grid Engine (SGE) cluster.

**Supplementary Note 3.7. Hard filtering of DPGT and GATK call sets.**

We followed GATK's documentation to do hard filtering of DPGT and GATK call sets.

VariantFiltration of GATK (version: 4.2.6.1) was used to mark variants that have not passed the filtering, while SelectVariants was used to apply filtering. The commands we ran was the following:

```
gatk VariantFiltration -V input.vcf.gz \
      --filter-expression "QD < 2.0 || FS > 60.0 || SOR > 3.0 ||
MQRankSum < -5.0 || ReadPosRankSum < -2.5 || ReadPosRankSum > 2.5" \
      -filter-name "HardFiltered" \
      --genotype-filter-expression "GQ < 5.0" \
      --genotype-filter-name "LowGQ" \
      --set-filtered-genotype-to-no-call \
      -O output.filter.vcf.gz

gatk SelectVariants --exclude-filtered \
      -V output.filter.vcf.gz \
      -O output.apply_filter.vcf.gz
```

**Supplementary Note 3.8. Softwares**

Software versions were DPGT version v1.2.8.0, Spark version 2.4.0-cdh6.2.0, Hadoop version 3.0.0-cdh6.2.0, GLnexus version 1.4.1, GATK version 4.2.6.1, sentieon versions sentieon-genomics-201911 and sentieon-genomics-202010.01, RTG tools version 3.12.1, bcftools version 1.13, bwa version 0.7.17-r1188, vcftools version 0.1.17.
